## Supplementary File 3 for "Improved base editing and functional screening in *Leishmania* via co-expression of the AsCas12a ultra variant, a T7 RNA Polymerase, and a cytosine base editor"

Julius-Maximilians-
UNIVERSITÄT
WÜRZBURG
:::::: Biozentrum
Zell- und Entwicklungsbiologie
.ZOO

LeishBASEedit

www.biozentrum.uni-wuerzburg.de/zeb/team/staff-scientists/tom-beneke/

**Dr. Tom Beneke**

Cell and Developmental Biology
Biocenter, University of Wuerzburg
Am Hubland • D-97074 Würzburg


### Cas12a-mediated integration of CBE sgRNA expression cassettes into *Leishmania*
### Protocol version February 2024

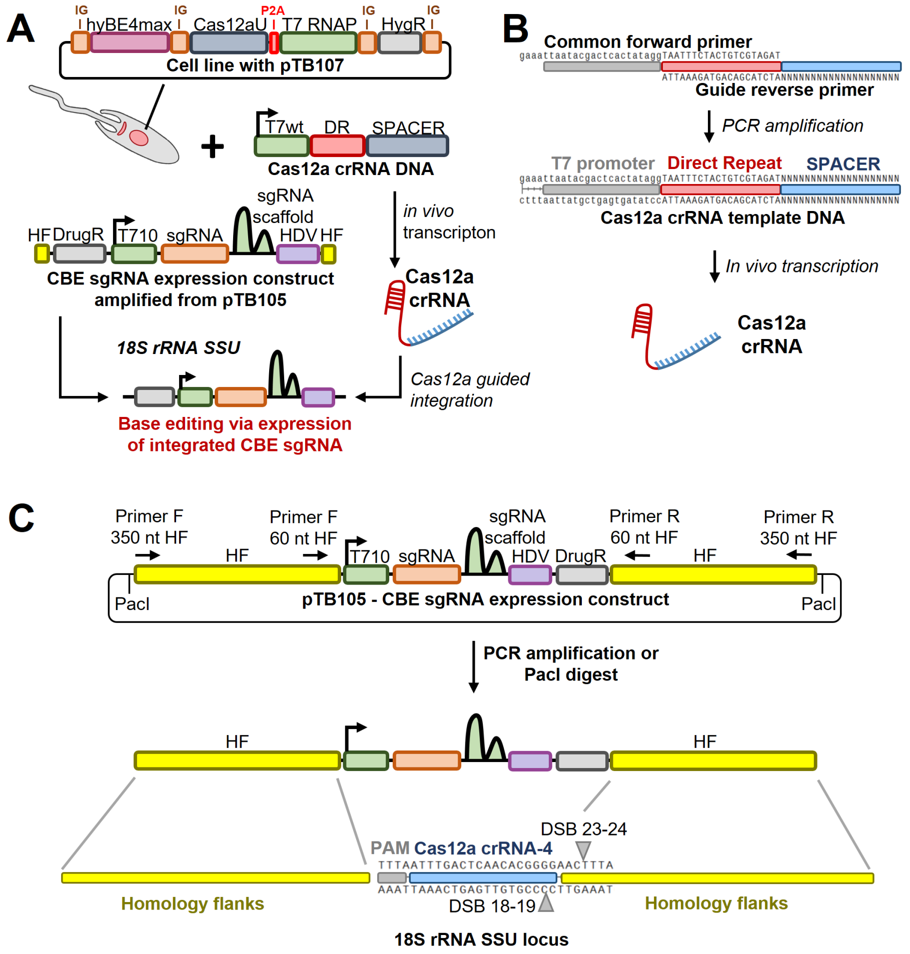


**Workflow overview.** The approach for Cas12a-mediated integration of the CBE sgRNA expression construct into the safe harbour 18S rRNA SSU locus is shown. **(A)** The pTB107 cell line expressing a Cas9 base editor (CBE), a Cas12a nuclease and a T7 RNAP is co-transfected with two constructs: (1) a Cas12a crRNA template DNA that can be *in vivo* transcribed by T7 RNAP to produce Cas12a crRNA (consisting of an unmodified T7 RNAP promoter [light green], a Cas12a direct repeat [DR, red] and a 20nt Cas12a guide target sequence [SPACER, blue]) and (2) a CBE sgRNA expression cassette that is integrated into the 18S rRNA SSU locus following the Cas12a-mediated double-strand break (cassette consists of two homology flanks [HF, yellow], a T7 T-10 GG promoter [light green], a sgRNA target sequence [orange], a Cas9 scaffold [dark green], a hammerhead ribozyme [HDV, purple] and a drug resistance marker [grey]). **(B)** The PCR amplification of the Cas12a crRNA template DNA is shown. A common forward and a guide specific reverse primer are used for amplification. The resulting amplicon can be used for *in vivo* transcription as described in (A). **(C)** The CBE sgRNA expression construct is generated by PCR or by PacI digest. Plasmid pTB105 contains 350nt homology flanks and therefore, the length of homology flanks can be varied when PCR amplifying the donor DNA cassette (e.g. 60 or 350nt homology flank). PacI digest can generate constructs with 350nt homology flanks only. The amplified PCR product is then integrated into the 18S rRNA SSU locus, following the Cas12a-mediated double strand break. The Cas12a SPACER (guide), PAM and homology flank sequences are indicated.

#### Step-by-step protocol

To integrate base editor guide expression cassettes into the 18S rRNA SSU locus, three main steps are required:

1. Cloning the CBE sgRNA into BbsI sites (see LeishBASEedit "Guide Cloning Protocol")
2. PCR amplify Cas12a crRNA template DNA and CBE sgRNA expression cassette
3. Transfection of PCR amplicons (see LeishBASEedit "Transfection Protocol")

This protocol describes Step 2.

**Primer preparation**

1. Prepare 100 µM stocks of the following standard desalted oligos (25nmol scale):
   - Cas12a crRNA template DNA primers:
     i. Common forward primer:
        5' gaaattaatacgactcactataggTAATTTCTACTGTCGTAGAT 3'
     ii. Guide specific reverse primer for integration into 18S rRNA SSU locus:
        5' TTCCCCGTGTTGAGTCAAATATCTACGACAGTAGAAATTA 3'
   - CBE sgRNA expression cassette primers:
     i. Forward primer for pTB105 amplification – 60nt homology flanks:
        5' GTTCGCAAGAGTGAAACTTAAAGAAATTG 3'
     ii. Forward primer for pTB105 amplification – 350nt homology flanks:
        5' GACCGCACCAAGACGAACTACAG 3'
     iii. Reverse primer for pTB105 amplification – 60nt homology flanks:
        5' CATCCTGTCCGGATCTGGTAAAG 3'
     iv. Reverse primer for pTB105 amplification – 350nt homology flanks:
        5' GTTCTCACTGACATTGTAGTGCGCG 3'

**PCR procotol**

For Cas12a crRNA DNA amplification prepare the following reaction mix (scale-up as needed):

| Reagent | Stock conc. | Final conc. | Volume per reaction |
|---|---|---|---|
| $ddH_2O$ | | | 35.00 µl |
| Phusion GC buffer | 5x | 1x | 10.00 µl |
| dNTP | 10 mM | 200 µM | 1.00 µl |
| Forward Primer | 100 µM | 2.0 µM | 1.00 µl |
| Reverse Primer | 100 µM | 2.0 µM | 1.00 µl |
| DMSO | 100% | 3% | 1.50 µl |
| Phusion™ High-Fidelity DNA Polymerase (F530S, Thermo Scientific™) | 2 unit/µl | 1 unit/50µl | 0.50 µl |
| | | | Total 50.00 µl |

Run reactions using the following cycle settings (a Hot Start is recommended):
Step 1: First denaturation 98°C 30 sec
Step 2: 98°C 10 sec
Step 3: 65°C 10 sec
Step 4: 72°C 10 sec
Step 5: 35x back to Step 2
Step 6: 72°C 7:00 min
Step 7: 4°C ∞
Step 8: End

For CBE sgRNA cassette amplification prepare the following reaction mix (scale-up as needed):

| Reagent | Stock conc. | Final conc. | Volume per reaction |
|---|---|---|---|
| $ddH_2O$ | | | 35.50 µl |
| Phusion GC buffer | 5x | 1x | 10.00 µl |
| dNTP | 10 mM | 200 µM | 1.00 µl |
| Forward Primer | 100 µM | 0.5 µM | 0.25 µl |
| Reverse Primer | 100 µM | 0.5 µM | 0.25 µl |
| Template DNA (pTB105) | 10ng/µl | 10 ng | 1.00 µl |
| DMSO | 100% | 3% | 1.50 µl |
| Phusion™ High-Fidelity DNA Polymerase (F530S, Thermo Scientific™) | 2 unit/µl | 1 unit/50µl | 0.50 µl |
| | | | Total 50.00 µl |

Run reactions using the following cycle settings (a Hot Start is recommended):
Step 1: First denaturation 98°C 30 sec
Step 2: 98°C 10 sec
Step 3: 65°C 10 sec
Step 4: 72°C 40 sec
Step 5: **20x** back to Step 2 (over-amplification should be avoided!)
Step 6: 72°C 7:00 min
Step 7: 4°C ∞
Step 8: End

**Alternatively, a PacI digest of 10 – 20 µg pTB105 plasmid in 50 µl can be performed for donor DNA preparation. But it will depend on the species, whether gel extraction following the digest is required!**

#### Preparing transfections

1. Verify success of digest and/or PCRs by running 2 µl of each reaction on a DNA agarose gel:
   - Cas12a crRNA template DNA should be 64 bp
   - CBE sgRNA cassette with 60nt HFs should be 1238 bp
   - CBE sgRNA cassette with 350nt HFs should be 1855 bp

2. If successful, mix remaining 48 µl of both reactions (96 µl total) and add 10 µl of 3M sodium acetate and 320 µl of 100% EtOH
3. Mix thoroughly by vortexing
4. Precipitate at -80°C for 1 hour or better at -20°C overnight
5. Centrifuge at 17,000g 4°C for 30 min
6. Wash pellet once with 500 µl 80% EtOH and centrifuge at 17,000g 4°C for 15 min
7. Take ethanol out and air dry pellet for 10-20 min
8. Add 50 µl $ddH_2O$ and let it dissolve at RT for 10 min
9. If not using sterile conditions, heat sterilize final product at 95°C for 5 min
10. Transfect entire product as described in LeishBASEedit “Transfection Protocol”
