## Supplementary figures and images for "Improved base editing and functional screening in *Leishmania* via co-expression of the AsCas12a ultra variant, a T7 RNA Polymerase, and a cytosine base editor"

### Source data 1 - Raw DNA images of Figure 2 S1B and E

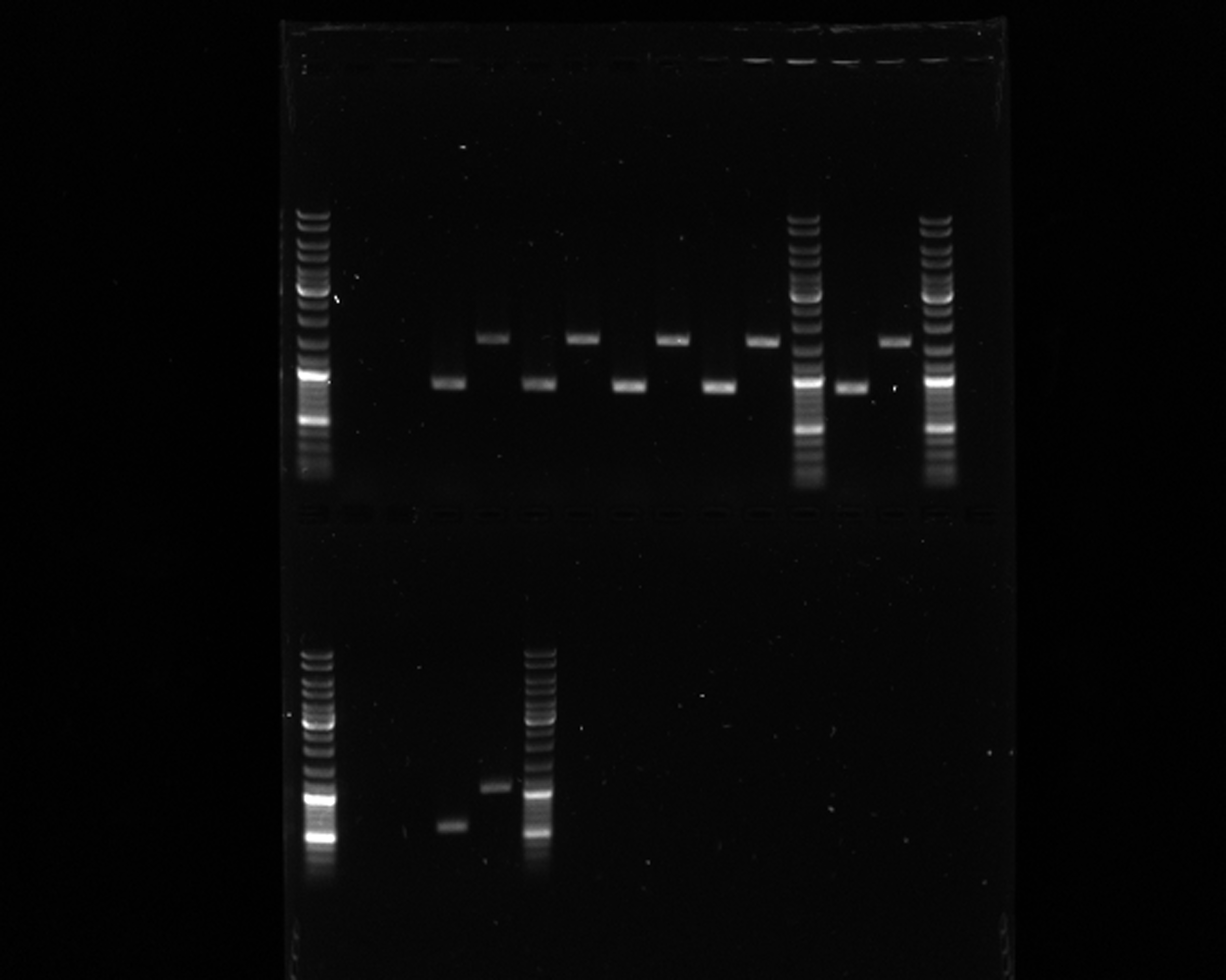

### Source data 2 - Raw DNA images of Figure 6B

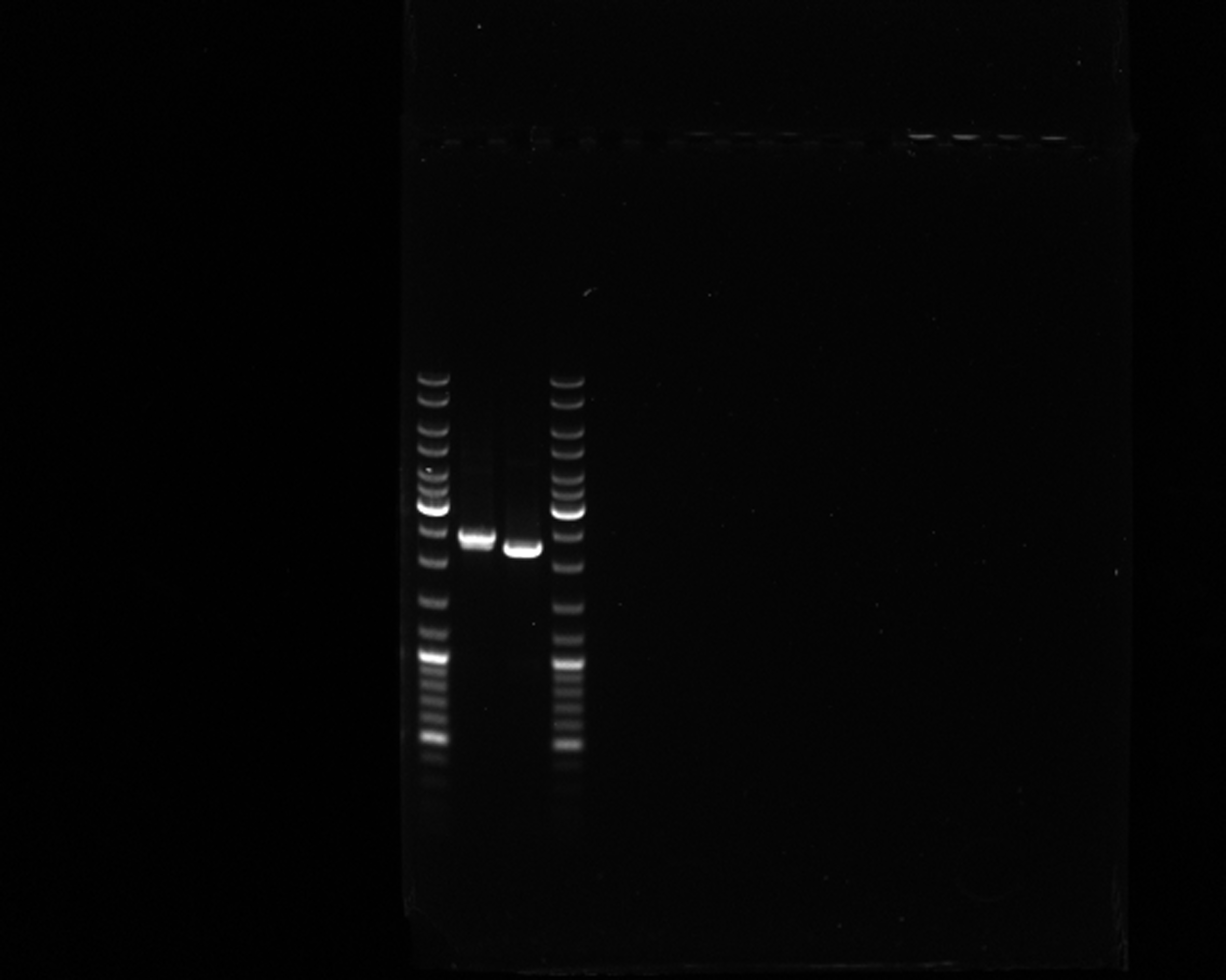
